# How to develop objective-driven comprehensive science outreach initiatives aiming at multiple audiences

**DOI:** 10.1101/023838

**Authors:** Sanjai Patel, Andreas Prokop

**Affiliations:** Faculty of Life Sciences, Michael Smith Building, Oxford Road, Manchester M13 9PT, UK

**Keywords:** *Drosophila*, science communication, outreach, public engagement, schools, university, teaching

## Abstract

Science outreach has become increasingly important for researchers and needs to be of ever improving quality, although the time available aside our science, teaching and administration activities is steadily decreasing. To square this circle, effective strategies are required. Here we argue that this can be achieved by setting simple but ambitious overarching objectives for comprehensive outreach initiatives which target multiple audiences, supported by cumulative build-up of shared high-quality resources, as well as the exchange and collaboration amongst scientists with a common outreach aim. To exemplify this strategy, we explain the low-budget, yet high-quality outreach initiative of the Manchester Fly Facility which aims to promote public awareness of the importance of the model organism *Drosophila* for biomedical research. (1) This initiative targets the general public at science fairs, through public videos, or through extracurricular activities in schools as well as the development of curriculum-relevant sample lessons for teachers - all supported by a dedicated website. (2) The initiative targets university students: by adapting the public outreach resources for their teaching, and through newly developed advanced training strategies that amalgamate the outreach objectives. (3) It targets fellow scientists through blogs, conference presentations and a second website that provides a one-stop-shop for resources, arguments and strategies. As will be explained, this multi-pronged approach is time-saving in the long run and it is powerful because it reaches a wide range of audiences, helps to gain momentum, to build resource, and to gradually improve quality through cross-fertilisation between different activities, and through exchange within the science community. This helps to build communities, and high-quality outreach will have further important added value: arguments that impress the public, tend to be most effective also with reviewers and grant panel members, and often help to readjust aspects of your own scientific work.

## Article

### Personal opportunities of public engagement activities

As basic scientists we have a vested interest in communicating our findings and concepts and use primarily publications, conferences and invited talks to communicate with specialist audiences and networks of colleagues, and increasingly also clinicians and industry to engage on the translational path. However, there is a strong urge to go a step further and engage in proactive science communication and outreach also with the general public. Budget cuts drive a political need to make the public and policy makers aware of the importance of our research, with a view to securing future funding. On the other hand, grant giving institutions expect us to explain the science financed by tax money, and to pay back to society in valuable currency: education (Rull, 2014). At first sight, this seems yet another task which adds to our already heavy duties in teaching, administration, group management, conference attendances, paper writing etc.

However, I would argue that public engagement activities can become a valuable time investment. Developing arguments fundamental enough so that they convince and excite non-specialist audiences is highly challenging, and it forces to think deeper and extract the essentials of ones science. This process is very likely to deliver new attitudes, scientific perspectives, ideas and strategies. Importantly, formulating arguments that grip members of the public tend to be particularly powerful and will appeal likewise to reviewers and members of grant panels.

### The importance of objective-setting

However, to achieve such added value, outreach needs to be of high quality. Self-evidently, it requires honest passion for the communicated science topic to be able to develop the necessary stamina and infect target audiences. In addition, it should be driven by clear long-term objectives.

A powerful example of effective objective setting was the Rhine Action Program^[1]^ agreed amongst the Rhine countries after the environmental disaster caused by the Santoz chemical spill in 1987. The key target for this program was to bring back the long-lost (since 1930s), pollution-sensitive salmon into the Rhine by the year 2000. Already in 1997, the salmon was back and pollution was reduced to 50-100% depending on substance. Obviously, an easy-to-understand overarching objective has inspired the creative implementation of an effective cocktail of measures that would hardly have been triggered by more specific, technocratic target setting.

Similarly, long-term objective setting for science outreach activities has important strategic advantages. Firstly, it helps to gradually raise the quality and momentum of a campaign, resulting in greater impact. Secondly, it provides room to think big and establish multifaceted initiatives which reach a wider range of target audiences or address the same audiences in different ways. Thirdly, long-term objectives are more likely to lead to substantial online resources that may develop self-sustaining dynamics and reach impact beyond the activities one directly engages in. Fourthly, it generates exciting opportunities to exchange ideas and resources, or even to collaborate, with other scientists in the field, thus weaving new communication networks.

### Awarness of the different target audiences

The different target audiences to be addressed by objective-driven, multifaceted science communication projects include the general public, but also university students and scientist colleagues.

Communicating to the general public involves a broad spectrum of audiences. Lay members of the public are typically addressed by street, pub or art fusion events, science fairs, theatrical happenings, school visits, lay articles in the press or on blogs, dedicated web sites or online videos. A particularly important target group within, are teachers and schools: Teachers may become efficient multiplicators of science outreach objectives, and addressing pupils provides powerful opportunities to achieve understanding and awareness by young audiences. Especially pupils may tell about their lesson experiences to friends or at home, thus spreading the message even further. Furthermore, most policy makers are non-specialist audiences, more likely to be reached by arguments developed for public outreach. Finally, it is often overlooked that scientists and clinicians in academia or industry, who are not from ones specific science field (but still sit on grant panels that judge applications on related science), may appreciate fundamental explanations originally designed to reach lay audiences.

University students are a very special target audience in our direct influence zone. In their initial years they can be considered lay audiences but, gradually, they need to be addressed as potential future scientists and colleagues who can carry the ideas behind your outreach objectives into the next generation.

Communication to fellow scientists serves the very different purpose of raising awareness about ones campaign, exchanging ideas and resources, and recruiting comrades-in-arms. This helps in several ways. It enormously facilitates the job of those who are new to public engagement. It pools ideas and resources from different sources, leading to collective improvement of outreach quality and avoiding the need for re-inventing the wheel. However, outreach to fellow scientists depends on efficient communication channels within scientific communities.

### An example for comprehensive outreach strategy promoting Drosophila research

In the following, I will use the science communication and outreach activities by the Manchester Fly Facility as an example in which we have gathered valuable experiences for 5 years with an objective-driven, comprehensive campaign.

Our overarching aim is to promote awareness of the enormous importance of the fruit fly *Drosophila* for biomedical research. Clearly, genetic invertebrate model organisms, such as *Drosophila*, have been crucial pillars in the process of scientific discovery for over 100 years (Bellen et al., 2010; Brookes, 2001/2002; Kohler, 1994), and most older generations have learned about flies at school. In more recent times of speedy and exciting methodological innovations, it gradually fades from general awareness that invertebrate model organisms remain important, cost-effective and prolific tools in contemporary biomedical research^[2]^. Students newly joining universities tend to hardly know about it, and it has recently been suggested that there is a decrease in public funding for *Drosophila* research (Wangler et al., 2015). As M. Brookes put it in his book about *Drosophila*: “*… we seem reluctant to accept that this tiny creature can teach us anything, let alone anything about ourselves”* (Brookes, 2001/2002).

Surprisingly, in spite of a long *Drosophila* research history, little concerted public outreach effort exists within the fairly large community of *Drosophila* researchers, likely because the fame of *Drosophila* in the past did not render this necessary ^[3]^. To promote the wider awareness of the importance of *Drosophila* research, we adopted therefore as our long-term objective, to raise the number of university students who know about the strategic importance of flies. As will become clear in the following, this simple and measurable, yet ambitious objective triggers activities that address a far larger range of target audiences than the actual objective might suggest at first sight.

### Targeting the general public at science fairs

One efficient way to raise awareness of *Drosophila* amongst students is to address young audiences before they enter university. This can be done very effectively in schools (see next section), but also in their family contexts, for example on public science events.

Public science fairs clearly are a challenge, since one has to shine amongst a multitude of other creative exhibition stands. One important strategy is to present high quality, catchy and self-explanatory visual displays which engage visitors. Some tricks for how they can be developed are explained elsewhere (Sumner and Prokop, 2013), and our various high-quality *Drosophila* displays are available upon request. In addition, we use a beamer to project a large-screen movie showing sequences of *Drosophila* tissue dynamics which are aesthetically very appealing and attract by-passers to stop and watch. Single sequences of this film display thought-provoking captions, such as “Wound healing” or “Moving blood cells”, animating visitors to engage in discussion (film available upon request).

A further helpful strategy is to organise various parallel displays/activities, each indicated by a big sign with a catchy title. This keeps visitors in the area and generates further opportunity to interact. We usually have stands with epileptic and paralytic flies (“Seizures and Paralysis”), for engaging in a climbing assay where visitors compare the motor performance of young versus old flies (“Testing Motor Skills”), for identifying genetic markers (“Learning to Fly”), for demonstrating optogenetic tools (“Fruit Fly Laser Quest”), and for microscopy of fluorescent tissues or larvae (“Seeing is Believing”; more detail available upon request).

Whilst younger audiences tend to engage in activities, their parents can be passive bystanders. Especially in the multi-display scenario, this provides opportunities for another strategy which targets primarily adults. For this, we have a central stand with the “Why the fruit fly?” poster which takes 5 minutes to explain and provides background information and exciting facts illustrating why and how *Drosophila* is used in research (avaliable upon request). Conjunct with this poster, we show film sequences comparing behaviours of flies and higher animals, such as boxing deer versus boxing flies, or a free-climber versus a blind flightless fly breeching across a gap of two stepping stones (available upon request). These and further examples (including alcohol addiction, learning, or circadian rhythm behaviours) are extremely effective in illustrating that properties expected to be typical human, rather reflect fundamental biology and can therefore be conceptually studied in flies ^[4]^. This central information stand is very effective in gaining the interest and understanding of teenagers or adults, thus strengthening the family experience.

Achieving a family experience enormously facilitates the “Trojan horse strategy” (Sumner and Prokop, 2013). This strategy encompasses that fliers are handed out which show display-related information or motives that children will find attractive to take home, such as Hama bead patterns of flies, fold plans of boxes with funny fly motives, games or riddles ^[5]^. If links to online resources are displayed on these fliers, this might tempt visitors to try them out at a later stage. Such online resources have to be suitable for lay audiences, designed to allow visitors to virtually re-live the event, find more detailed explanations and additional background information, as well as links inviting to further explore aspects around *Drosophila* in the world-wide web (see our online materials in the next section).

A further strategy to gain more impact is to join up with related outreach teams and form part of a larger theme. For example, we once successfully combined our *Drosophila* neurobiology show with displays by other groups on stroke and neurodegeneration in rodents, pig brain dissection sessions, and neurosurgeons displaying sham brain operations or activities where visitors cut open training skulls. On the one hand, the fly stands provided helpful information on fundamental neurobiology and increased the educational value of the event, on the other it facilitated our objective of illustrating how fly research contributes to the field.

Taken together, science fairs are a fantastic playground for outreach, since one receives immediate and telling responses from visitors. This provides opportunities to explore and test new ideas, and learn which arguments work best with lay audiences and how to package and optimise them. As an important strategy for constant improvement, it is helpful to organise de-briefing sessions within days after the event and set clear objectives for implementing quality-enhancing changes before the next occasion. This increases quality and impact, and it keeps the activities more lively and interesting also for the performers. Furthermore, science fair activities can have enormous added value by generating concepts and ideas which can be applied to other outreach activities. For example, the argumentation strategy we developed for the “Why the fruit fly?” poster, has delivered the basic framework for our two YouTube videos entitled “Small fly, big impact” ^[6]^, as well as for introductory presentations we often give at schools or on teacher seminars (available upon request). We also realised that most of the activities we developed for science fair stands are ideal for school demonstrations, thus generating important new opportunities to capitalise on the time invested in their development.

### Targeting schools

Our school work pursues at least two fundamental strategies: firstly, we visit schools for extra-curricular days; secondly, we develop and provide sample lessons that teachers can adopt independently for their biology teaching.

Our school visits reach about 200 pupils per day and require 4-5 helpers, some computer equipment, 20 low-cost microscopes, a hand-held fluorescent lamp and numerous smaller items ^[5]^ (detail available upon request). We give an introductory lesson followed by two rotation cycles of three to four parallel 25 minute demonstration sessions ^[7]^. These school events aim to achieve a number of important goals. (1) They aim to enthuse pupils and enhance their general interest in science and, perhaps, a university career. (2) They provide opportunities to promote the content and key messages of our outreach. (3) They help pupils in their curriculum which, in return, enhances the value and impact of these school events enormously.

Especially the last point about curriculum relevance is often overlooked and requires background knowledge of the actual biology specifications ^[8]^. In daily school routines, these biology specifications are often taught as isolated learning objectives with insufficient conceptual background and synopsis, and it is difficult for teachers to weave them into relevant contexts that reflect contemporary science. This is a great opportunity for scientists since they can use their expertises to provide conceptual backgrounds and explain relevant contemporary research contexts. It is a great opportunity for *Drosophila* since it is the animal with the conceptually best understood biology (ideal for teaching), it is actively used in research (ideal to convey relevance), and it is ideally suited for school experiments (ideal to make lessons more exciting and memorable).

To capitalise on these opportunities, we use PowerPoint presentations for each of our sessions, which explain curriculum-relevant conceptual backgrounds of the incorporated experiments, and how these experiments are applied in science (available upon request). For example, in our neurobiology lesson we explain fundamental principles of neuronal circuits, including action potentials and synapses, and combine this with experiments involving epileptic flies, neuronal silencing in temperature-sensitive *shibire* mutant flies as well as optogenetic tools, all accompanied by examples of their scientific applications.

To enhance the learning outcome even further, it is useful to include follow-up tasks, such as essay writing about the school science event ^[9]^, and teachers who arranged science days are usually supportive of such add-ons. The learning outcome of such tasks can be significantly improved if supportive resources, printed or online (see below), can be provided to students to facilitate and enhance their revision and performance on the task.

Our second strategy involves the development of curriculum-relevant sample lessons to be adopted by teachers, where *Drosophila* is used as a modern teaching tool, shoulder to shoulder with human examples. These lessons are accompanied by self-explanatory support materials so that teachers can implement and use them independently. To achieve this, we initiated the droso4schools project, where two PhD students were sent for one month as teaching assistants into two collaborating schools in order to gain valuable teaching experiences and discuss with teachers about strategies, requirements and suitable biology topics ^[10]^. Capitalising on these experiences, sample lessons were developed and subsequently tested at the schools, and they are now available for download together with adjunct materials ^[11]^. One lesson uses a fly climbing assay to teach statistics and concepts of ageing and neurodegeneration, another represents a synoptic approach using the topic of alcohol to cover about 7 biology specifications from gene concepts to enzymes and even evolution, and a third lesson introduces to classical genetics whilst explaining fundamental discovery processes in science. They all involve experiments with flies and provide exciting opportunities to discuss socially relevant topics.

Importantly, this droso4schools project is supported by a dedicated webpage which provides background information and summary pages for the uploaded lessons (e.g. ageing and neurodegeneration, statistics), a tab explaining the rationale for using *Drosophila* in research, and a tab comparing the organs of flies and humans ^[12]^. This website is ideal for lesson preparation or revision as well as for homework tasks.

### Targeting university students

At universities, new students can be considered lay audiences. Therefore, resources originally designed for science fairs and schools are very effective for introducing students to scientific concepts and strategies. We do this successfully using many of the above mentioned resources to introduce students to *Drosophila* as an important invertebrate model organism in biomedical research. In the next step, to train students up as future scientists, we developed and published further strategies and resources.

For example, our *Drosophila* training package provides self-study strategies to teach students at PhD level about the fundamentals of fly genetics and develop their skills to design complex genetic crosses (Roote and Prokop, 2013). An essential part of this package is the “Rough guide to *Drosophila* mating schemes” (Prokop, 2013a) which introduces to the specifics and practicalities of classical fly genetics and transgenic technologies. In addition, it provides a historical perspective and the rationale for using flies, all-in-all aiming to equip students with fundamental understanding of *Drosophila* as a model and, in turn, the ability to better explain the importance of their work to others.

To reach larger student numbers already at the undergraduate level, we stripped the rough guide down to a light version (Prokop, 2013b) and used it on second year university courses with up to 65 students. We demonstrated that the entire genetics training can be performed in this context and, to promote its use as an accredited course element, we developed an efficient hybrid strategy for the electronic assessment of virtually all aspects of the learned skills repertoire. As explained elsewhere ^[13]^ (Fostier et al., 2015), this assessed training represents active learning and a modern approach to convey core knowledge of classical genetics. Furthermore, it promotes essential understanding of *Drosophila* as a model organisms, and teaches problem solving skills as a desirable outcome of university education.

### Targeting fellow drosophilists

We argued in the second section of this article that objective-driven, multi-facetted outreach provides exciting opportunities to establish new collaborative and communication networks amongst scientists, eventually leading to enhanced quality and impact. This requires effective means of dissemination for achieving a wider awareness amongst the respective scientists about the outreach objectives as well as available strategies and resources.

The *Drosophila* community had a long tradition of excellent information exchange and dissemination (Kelty, 2012), but ironically lost this advantage when entering the era of the internet and social media. Therefore, to enhance awareness of *Drosophila* outreach, we write blogs ^[13, 14]^, present on conferences ^[15]^, and developed and curate the “For the public” area on the Manchester Fly Facility website ^[16]^, which represents a one-stop-shop for links and resources concerning *Drosophila* outreach, including large collections of educational resources, lay and history articles about fruit fly research, and an embedded feed of relevant *Drosophila*-specific tweets ^[17]^ for those who are not on Twitter. This site is complementary to our droso4schools as well as other *Drosophila*-specific webpages with important science communication content (e.g. FlyBase, FlyMove, Interactive Fly, Fly on the Wall), and it constitutes therefore a new and helpful resource.

### Concluding thoughts

The examples described in this article were implemented with fairly low amounts of dedicated funding and mainly driven by the two authors further supported by many active and creative contributions of student and postdoctoral volunteers. We hope they illustrate how overarching objectives can become an important and powerful driver for multi-facetted outreach activities which target a wide spectrum of audiences. Notably, such strategies help to improve quality of outreach, through providing opportunities for gradual improvements over time, through communication and exchange within science communities, and through cross-fertilisation between different activities within ones own outreach campaign. Thus, as illustrated by a few examples in this article, the engagement in one activity often triggers ideas that are similarly powerful and applicable in other activities, and this offers excellent opportunities for quality enhancement across ones campaign.

A key goal of outreach is to achieve impact. For this, substantial resource of high quality needs to be developed and then effectively disseminated. A great number of our resources were described here, but many more strategies can be tried ^[18]^, such as longer-term school projects (e.g. in afternoon clubs) to perform genetic screens, to experiment with population genetics (e.g. enrich fly populations for alcohol tolerance), or to study fly behaviours and couple this with programming skills capitalising on the advantages of low-cost computing power (e.g. Raspberry Pi). Regarding the importance of dissemination, we have described here how we address different target audiences, but key challenges remain. For example, we need to disseminate our resources more widely amongst school teachers. To achieve this, we have chosen, as our first strategy, to write manuscripts for teacher and student journals which, once published, will be used as key references for targeting teacher associations and for presenting on teacher seminars and conferences. As a further example, policy makers will have to be targeted. Whilst this will be easier to achieve with the lobbying capacity of learned societies, the arguments and resources developed through public outreach will form a very helpful foundation to this end.

Ultimately, outreach needs to lead to measurable impact, and the evaluation procedures and strategies through which such impact can be documented need to be carefully considered, planned and implemented ^[19]^. There is never a guarantee that science outreach will have a measurable outcome, and this risk is the flip side of any public engagement. However, we hope that some of the ideas and strategies provided in this article will help to minimise these risks and enhance the success of any public outreach you may be engaged in.

## Acknowledgements

We would like to thank the many students and postdocs of the Manchester fly community who have helped over the years to implement and sustain the various outreach activities. AP is supported by BBSRC project grants (BB/I002448/1, BB/L000717/1 and BB/M007553/1), and SP through various BBSRC and MRC grants.

## Footnotes

1 www.bbc.co.uk/scotland/education/int/geog/eei/rivers/rhine/strategies/info.shtml?strategies=1

2 droso4schools.wordpress.com/why-fly

3 blog post: poppi62.wordpress.com/2015/05/12/fly-scicomm

4 see some examples here: droso4schools.wordpress.com/alcohol/#7

5 see some examples here: www.flyfacility.ls.manchester.ac.uk/forthepublic/outreachresources/#Equipment

6 droso4schools.wordpress.com/why-fly/#Movies

7 Schedule for and essays about a school visit: flyfacility.files.wordpress.com/2014/11/cheadlehulmeessays.pdf

8 find information under the first link of this page: bsdb.org/public-outreach

9 Schedule and essays of a school visits: flyfacility.files.wordpress.com/2014/11/cheadlehulmeessays.pdf

10 see our YouTube video: www.youtube.com/watch?v=DQKFtt3p2C8

11 figshare.com/articles/Biology_lessons_for_schools_using_the_fruit_fly_Drosophila/1352064

12 droso4schools.wordpress.com

13 blog post: poppi62.wordpress.com/2015/05/12/fly-scicomm

14 blogs.brandeis.edu/flyonthewall/why-do-we-have-to-learn-this-stuff-establishing-drosophila-as-a-modern-teaching-tool-in-schools

15 files.figshare.com/1939326/15_03_07_ADRC_workshop_Prokop.pdf

16 www.flyfacility.ls.manchester.ac.uk/forthepublic/teachersandschools

17 @interactivefly, @fly_papers, @FlyBaseDotOrg

18 *Drosophila*-specific ideas: www.flyfacility.ls.manchester.ac.uk/forthepublic/outreachresources/#Teaching; general ideas: bsdb.org/public-outreach

19 www.publicengagement.ac.uk/plan-it/evaluating-public-engagement & www.manchesterbeacon.org/files/manchester-beacon-pe-evaluation-guide.pdf & www.ucl.ac.uk/public-engagement/evaluation

## References

Bellen, H.J., Tong, C., Tsuda, H., 2010. 100 years of *Drosophila* research and its impact on vertebrate neuroscience: a history lesson for the future. Nat Rev Neurosci. 11, 514–522.

Brookes, M., 2001/2002. Fly: The Unsung Hero of Twentieth-Century Science, Vol., Ecco/Phoenix.

Fostier, M., Patel, S., Clarke, S., Prokop, A., 2015. A novel electronic assessment strategy to support applied *Drosophila* genetics training on university courses. G3 (Bethesda). 5, 689–98.

Kelty, C.M., 2012. This is not an article: Model organism newsletters and the question of open science’. BioSocieties. 7, 140–68.

Kohler, R.E., 1994. Lords of the fly. Drosophila genetics and the experimental life, Vol., The University of Chicago Press, Chicago, London.

Prokop, A., 2013a. A rough guide to Drosophila mating schemes (in: Roote, J and Prokop, A., 2013, “How to design a genetic mating scheme: a basic training package for Drosophila genetics” G3/Bethesda 3, 353–8). http://dx.doi.org/10.6084/m9.figshare.106631.

Prokop, A., 2013b. 2^nd^ year *Drosophila* developmental genetics practical. http://dx.doi.org/10.6084/m9.figshare.156395

Roote, J., Prokop, A., 2013. How to design a genetic mating scheme: a basic training package for *Drosophila* genetics. G3 (Bethesda). 3, 353–8.

Rull, V., 2014. The most important application of science. EMBO Rep. 15, 919–922.

Sumner, J., Prokop, A., 2013. Informing the general public about cell migration - an outreach resource. http://dx.doi.org/10.6084/m9.figshare.741264

Wangler, M.F., Yamamoto, S., Bellen, H.J., 2015. Fruit flies in biomedical research. Genetics. 199, 639–53

